# Estradiol-induced progesterone synthesis develops post-puberty in the rostral hypothalamus and coincides with post-pubertal changes in the steroidogenic pathway in female mouse hypothalamic astrocytes

**DOI:** 10.1101/833434

**Authors:** M.A. Mohr, T. Keshishian, B.A. Falcy, B.J. Laham, AM Wong, P.E Micevych

## Abstract

The development of estrogen positive feedback is a hallmark of female puberty. Both estrogen and progesterone signaling are required for the functioning of this neuroendocrine feedback loop but the physiological changes that underlie the emergence of positive feedback remain unknown. Only after puberty does estradiol (E2) facilitate progesterone synthesis in the rat female hypothalamus (neuroP), an event critical for positive feedback and the LH surge. We hypothesize that prior to puberty, these astrocytes have low levels of membrane estrogen receptor alpha (ERα), which is needed for facilitation of neuroP synthesis. Thus, we hypothesized that prepubertal astrocytes are unable to respond to E2 with increased neuroP synthesis due a lack of membrane ERα. To test this, hypothalamic tissues and enriched primary hypothalamic astrocyte cultures were acquired from pre-pubertal (postnatal week 3) and post-pubertal (week 8) female mice. E2-facilitated progesterone was measured in the hypothalamus pre- and post-puberty, and hypothalamic astrocyte responses were measured after treatment with E2. Prior to puberty, E2-facilitated progesterone synthesis did not occur in the hypothalamus, and mERα expression was low in hypothalamic astrocytes, but E2-facilitated progesterone synthesis in the rostral hypothalamus and mERα expression increased post-puberty. The increase in mERα expression in hypothalamic astrocytes corresponded with an increase in caveolin-1 protein, PKA phosphorylation, and a more rapid [Ca^2+^]_i_ flux in response to E2. Together, results from the present study indicate that E2-facilitated neuroP synthesis occurs in the rostral hypothalamus, develops during puberty, and corresponds to a post-pubertal increase in mERα levels in hypothalamic astrocytes.

**SIGNIFICANCE STATEMENT:** Estradiol facilitation of hypothalamic neuroprogesterone synthesis is necessary for the positive feedback of the LH surge. The present study localized the increase of neuroprogesterone to the rostral hypothalamus, a region that mediates estrogen positive feedback. Across pubertal development, hypothalamic astrocytes increase levels of membrane ERα and the cell signaling responses needed to facilitate neuroprogesterone synthesis that triggers the LH surge demonstrating a mechanism for pubertal maturation resulting in reproductive competence.

## INTRODUCTION

Puberty is a complex developmental stage that elicits a cascade of neuroendocrine changes leading to reproductive maturation. In females, the development of reproductive circuits during puberty culminates in the brain responding positively to ovarian estradiol (E2), triggering the luteinizing hormone (LH) surge that underlies ovulation. There is mounting evidence that alongside E2, brain-derived progesterone (neuroP) is needed for full activation of estrogen positive feedback: 1) astrocyte-derived neuroP is crucial for fully stimulating kisspeptin release in an in vitro model of adult anterior hypothalamic kisspeptin neurons (1, 2), 2) blocking neuroP synthesis arrests estrous cyclicity and decreases the LH surge (3, 4), and 3) nuclear progesterone receptor (PGR) signaling in anteroventral periventricular (AVPV) kisspeptin neurons is required for the LH surge and fertility in mice (5); Mohr et al., in review). However, whether neuroP synthesis develops during puberty in the female mouse remains unknown.

Steroid hormones, including E2 and progesterone (P4), act directly in the cell nucleus as well as at the membrane, initiating non-classical signaling that elicits intracellular signaling that may also affect transcription (6), E2 membrane-initiated signaling elevates intracellular calcium inducing rapid neuroP synthesis (7). E2-signaling in astrocytes is initiated via membrane-associated ER-alpha (mERα) transactivation of metabotropic glutamate receptor 1a (mGluR1a; (8)). The shuttling of mERα to the plasma membrane and subsequent coupling to mGluR is regulated by a class of lipid raft proteins, called caveolins (9). In a process necessary for neuroP synthesis, caveolin-1 shuttles full-length mERα to the plasma membrane in association with mGluR1a. E2 binding initiates excitatory downstream signaling cascades, including IP_3_ receptor-dependent intracellular Ca^2+^ flux (10) and the phosphorylation of PKA (11), which in turn activate steroidogenic acute regulatory protein (StAR) and translocator protein (TSPO). These proteins mediate the transport of cholesterol into the mitochondrial matrix where CYP11A1 cleaves the side chain forming pregnenolone. Finally, 3β-hydroxysteroid dehydrogenase (3β- HSD) converts pregnenolone into P4 in the smooth endoplasmic reticulum.

Like the LH surge, E2-facilitated neuroP synthesis seems to occur solely in post-pubertal female rodents. Indeed, astrocytes derived from the adult female rodent hypothalamus respond to E2 stimulation with an increase in neuroP synthesis, and hypothalamic astrocytes harvested from neonatal female rodents or male rodents did not appear to respond to E2 with increased neuroP synthesis (12). Recently, we (Mohr et al. 2019), formally tested whether systemic administration of E2 to pre- and post-pubertal ovariectomized rats induced neuroP in the hypothalamus. Only post-pubertal female rats had elevated hypothalamic neuroP levels. Moreover, neuroP levels were elevated on proestrus in gonadally intact female rats. Thus, *in vivo* and *in vitro* studies suggest that during puberty, hypothalamic astrocytes develop the cellular machinery to have E2-induced neuroP synthesis. To explore the hypothesis that the E2- faciliation of neuroP occurs during puberty, the present study examined the neuroanatomical location and development of neuroP synthesis *in vivo*, and the sequelae of E2 action in astrocytes that are associated with neuroP synthesis: E2-induced [Ca^2+^]_i_, levels of membrane ERα and expression of proteins associated with steroidogenesis pre- (postnatal week 3) and post-pubertal (postnatal week 8) development.

## MATERIALS AND METHODS

All experimental procedures were performed in accordance with the principles and procedures of the National Institutes of Health Guide for the Care and Use of Laboratory Animals and approved by the Chancellor’s Animal Research Committee at the University of California, Los Angeles (Protocol Number: ARC # 2000-077).

### Animals

All C57Bl/6 female mice were purchased from The Jackson Laboratory (Bar Harbor, ME). Prepubertal mice for the *in vivo* neuroP experiment were shipped to arrive at PND10 with their dams. For the post-pubertal *in vivo* neuroP group, mice arrived on PND 35. Prepubertal mice used for the primary astrocyte cultures were shipped post-weaning (PND 21) and arrived on PND23. Upon arrival, mice were housed 4 per cage in a climate-controlled room, with a 12-hr light, 12-hr dark cycle. Food and water were available ad libitum to the mice.

### OVX and E2 treatment to elicit neuroP

An estradiol paradigm that mimics both estrogen positive and negative feedback was used to elicit neuroP [Dror, 2013]. One week after arrival, female mice were bilaterally OVX and implanted with a subcutaneous (s.c.) pellet containing 1 µg 17-beta-estradiol (EB) in safflower oil (to mimic estrogen negative feedback). Five days later, mice were injected with 1 µg/20 g BW (s.c.) EB dissolved in safflower oil (to mimic positive feedback), or oil vehicle containing only safflower oil. Brains were collected the next day, one hour prior to lights-off, when prepubertal mice were PND 23, and post-pubertal mice were PND 48.

### Hypothalamic dissections to measure neuroP

Mice were deeply anesthetized with isoflurane, then decapitated and brains were removed. A mouse brain matrix and razor blades were used to obtain hypothalamic blocks using the following boundaries: rostral extent of the optic chiasm, rostral extent of the mammillary bodies, lateral edges of the tuber cinereum, and the top of the third ventricle. 2 mm of rostral hypothalamus and 3 mm of mediobasal hypothalamus were collected and weighed (see Figure 1). Rostral hypothalami and mediobasal hypothalami samples were homogenized using a Dounce homogenizer in 500 µl and 1000 µl of PBS respectively. Samples were then sonicated and stored at -80° C until extraction.

**Figure 1.**
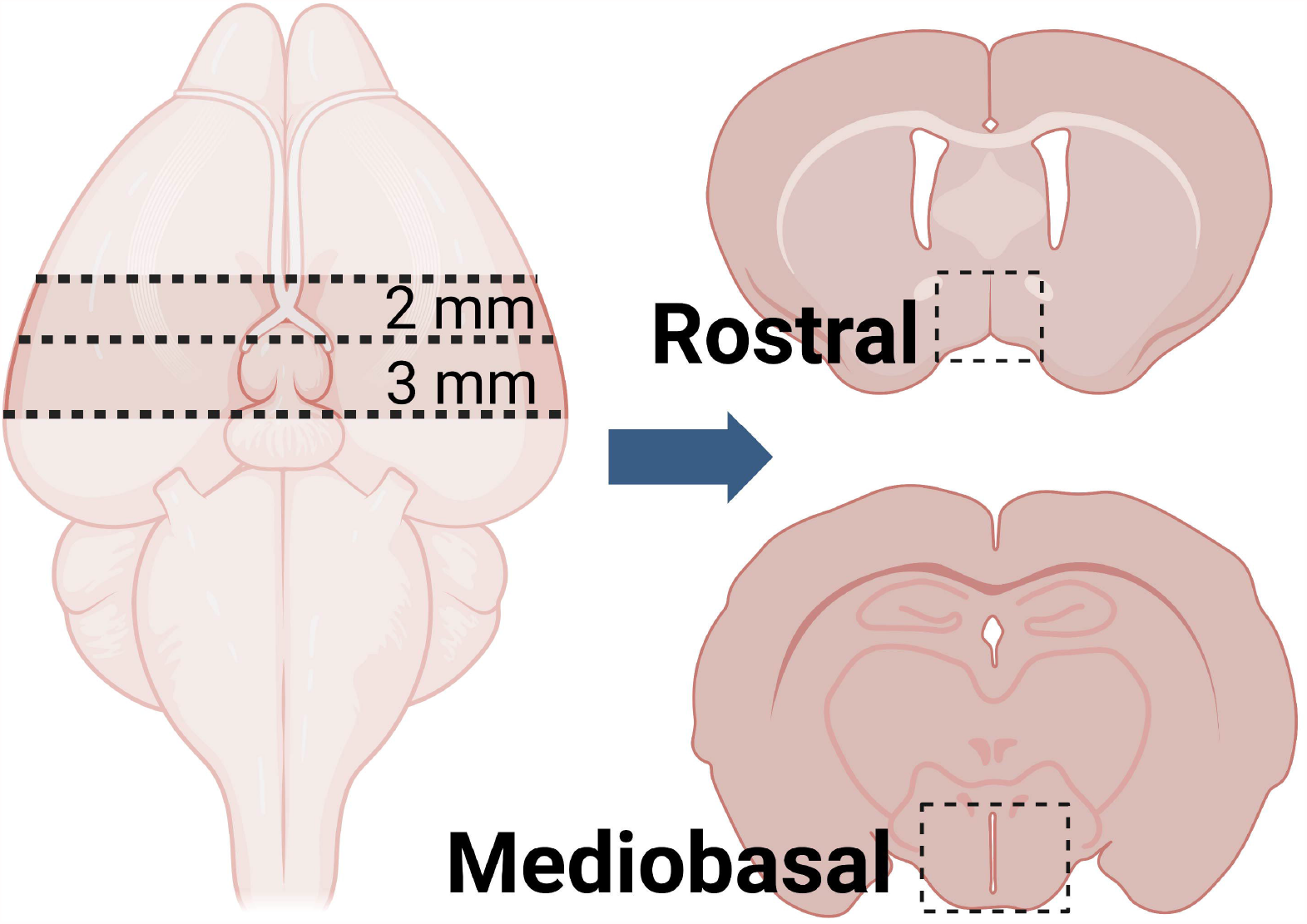
Location of hypothalamic dissections. To measure E2-induced neuroprogesterone, rostral and mediobasal hypothalamic blocks were taken using the boundaries indicated by dotted lines.

### Progesterone Extraction and ELISA

Progesterone was extracted from frozen samples using diethyl ether. Briefly, an equal volume of diethyl ether was added to homogenized tissue and vortexed until aqueous and lipid layers were separated. Bottom aqueous layer was discarded. The remaining lipid layer containing steroids was dried under nitrogen gas and evaporated samples were stored at -20° C until analysis. Frozen samples were reconstituted in assay buffer (Cat# 80-0010) before analysis. Enzo Scientific Progesterone ELISA kit (cat #ADI-900-011) was used to measure [P4] per manufacturer’s instructions. However, the standard curve was modified to include concentrations between 0.977-250 pg/ml.

### Primary astrocyte culture

Primary hypothalamic astrocyte cultures were prepared from a total of 65 C57Bl/6 female mice (Jackson) consisting of 2 ages: postnatal day (PND) 23 (n = 34) and 60 (n = 29). Cultures were generated from cohorts containing 8 animals (n = 4/age) at a time. Hypothalamic blocks were dissected with the following boundaries: rostral extent of the optic chiasm, rostral extent of the mammillary bodies, lateral edges of the tuber cinereum, and the top of the third ventricle. The tissue was minced, dissociated by enzymatic treatment with 2.5% trypsin solution (Invitrogen) with DNase I (Sigma #D4263) and incubated in a 37°C water bath where it was agitated every 10 min for 30 min. The trypsin was removed and 5 ml DMEM/F12 (Mediatech), supplemented with 10% fetal bovine serum (FBS) (Invitrogen), 1% penicillin (10000 IU/mL)/streptomycin (10000 μg/mL) solution (Mediatech), and 2 mM L-glutamine (Hyclone) was added. The tissue was triturated 6 times with a 5 ml pipette. Cells were plated in T-75 flasks with 10 ml DMEM/F12 (Mediatech), supplemented with 20% fetal bovine serum (FBS) (Invitrogen), 1% penicillin (10000 IU/mL)/streptomycin (10000 μg/mL) solution (Mediatech), and 2 mM L-glutamine (Hyclone) and maintained at 37°C with 5% CO2 for 3 days before the medium was replaced. After 1 week, FBS concentration was decreased to 15%. After 2 weeks, FBS concentration was decreased to 10%. Astrocyte cultures were grown to confluence and shaken on an orbital shaker at 200 rpm for 4 hours to eliminate oligodendrocytes and microglia, yielding cultures with >95% glial fibrillary acid protein-immunoreactive astrocytes. Before experimental manipulations, astrocytes were steroid- starved for 2 hours with 5% charcoal-stripped dextran-treated FBS in phenol-free DMEM/F12 medium. Cultures were then treated with vehicle dimethylsulfoxide (DMSO; Sigma-Aldrich, cat# D2650) or 17β-estradiol (1 or 10 nM; Sigma-Aldrich, cat #: E-8875) in steroid-free media (see above) for 1 hour at 37°C. After E2 treatment cells were collected for immunoblotting.

### Calcium Imaging

Astrocytes (375,000-570,000) were plated onto poly-d-lysine (0.1 mg/ml; Sigma-Aldrich)-coated glass coverslips and 1-2 days later pretreated for 2 h with 5% charcoal-stripped FBS DMEM/F- 12 medium. Before imaging, cells were loaded with the calcium indicator Fluo 4-AM (3 μM; Thermo-Fisher; F14201) dissolved in HBSS with 0.3% DMSO and 0.06% Pluronic F-127 (Thermo-Fisher; P3000MP) for 45 min at 37°C. HBSS was used to wash the cells and then glass coverslips were mounted onto a 50-mm chamber insert (Warner Instruments) fixed into 60 × 15-mm cell culture dish (Dow Corning) and placed into a QE-2 quick exchange platform (Warner Instruments) for imaging on a Zeiss Examiner D1 microscope, using an Achroplan 20/0,50 water immersion objective (Zeiss). A perfusion pump (1 ml/min, 37°C) was used to deliver HBSS before and after application of E2 (100 pM diluted from DMSO-stocks in HBSS, 17β-estradiol, Sigma-Aldrich, cat # E-8875) or ATP (30 uM diluted in HBSS; Sigma Aldrich, cat# A2383). Fluo-4AM fluorescence was excited with the 488-nm argon laser, and emission signals above 505 nm were collected. Videos consisted of 1200 frames (512 × 512) taken, on average, every 161.08 ms (6.2 Hz, +/- 0.4 ms). Calcium flux was measured as delta F/F, or the change in fluorescence intensity after E2 or ATP treatment over the fluorescent intensity during HBSS treatment before stimulation. Peak values were defined as the highest delta F/F value obtained after the perfusion with E2 or ATP. Responding cells were those whose peak values were at least 20% above the baseline value prior to E2 or ATP delivery.

### Cell Surface Biotinylation

Astrocytes were pretreated for 2 h with 5% charcoal-stripped FBS DMEM/F-12 medium and incubated with DMSO vehicle or in the presence 1 nM 17-estradiol (Sigma-Aldrich) in 5% charcoal-stripped FBS DMEM/F-12 medium for 1 h at 37°C. Cells were then washed three times with ice-cold HBSS with Ca^2+^ and Mg^2+^ buffer and incubated with freshly prepared 0.5 mg/ml EZ-Link Sulfo-NHS-LC-Biotin (Pierce) in HBSS at 4°C for 30 min with gentle agitation. Excess biotin reagent was quenched by rinsing the cells three times with ice-cold glycine buffer (50 mM glycine in HBSS). Cells were scraped into 10 ml of HBSS solution, transferred into a 50 ml conical tube, and centrifuged at 850 × g for 5 min. The pellet was resuspended in 200 μl of RIPA Lysis Buffer (Santa Cruz Biotechnology) containing the following proteases inhibitors: 1 mM phenylmethylsulfonyl fluoride, 1 mM EDTA, 1 g/ml pepstatin, 1 g/ml leupeptin, 1 g/ml aprotinin, and 1 mM sodium orthovanadate (all inhibitors were from Santa Cruz Biotechnology). The cells were homogenized by passing through a 25-gauge needle 10 times every 10 min for a total of 30 min. The cell extract was centrifuged at 14,000 rpm for 5 min at 4°C, and the protein concentration of the supernatant was determined using the bicinchoninic acid (BCA) Assay (Pierce). Two hundred microliters of each sample with 1500 g/ml protein concentration was incubated with 200 μl of Immobilized NeutrAvidin Gel (Pierce) overnight at 4°C and spun for 1 min at 1000 × g. The beads were washed four times with 500 μl of radioimmunoprecipitation assay (RIPA) buffer (Santa Cruz Biotechnology) containing the same proteases inhibitors mentioned previously. The bound proteins were eluted with Laemmli Buffer containing BME (950 μl Laemmli, 50 μl of BME) for 2 h at 37°C before centrifugation at 14000 rpm for 5 min.

### Immunoblotting

For Western blotting, astrocyte cultures were washed three times with 10 mL PBS buffer and collected by mechanical scraping (Corning). The samples were centrifuged (850 × g for 5 minutes), aspirated of PBS, and lysed in RIPA lysis buffer (Santa Cruz) with protease inhibitor and phosphatase inhibitor (Invitrogen). Pellets were sonicated at 3 × 10 pulses and kept on ice for 30 min before 4°C centrifugation at 14000 × g for 5 min. Protein concentrations were determined using the BCA Assay (Pierce). Protein (10 μg of whole-cell lysates; 40 μl of surface biotinylated lysates) of each sample was loaded on an 8-16% (for TSPO, 3b-HSD, and StAR) or 4-20% (for mERα, caveolin-1, and PKA) SDS precast gels (Bio-Rad) and separated by electrophoresis. Proteins were transferred onto PVDF (GE) membranes using electrophoresis at 11 V for 18 hours (Invitrogen). Because GAPDH protein changed across puberty (data not shown) a Ponceau S (0.5% Ponceau S with 1% acetic acid) stain was done for westerns using whole-cell lysates and the 45 kDa band was imaged and used as a loading control (13). Membranes were probed with primary rabbit or mouse monoclonal or polyclonal antibodies diluted in 5% bovine serum albumin or nonfat dry milk in Tris-buffered saline/Tween 20 (0.1% [vol/vol]) according to the suppliers’ recommendations (see Table 1). Primary incubations lasted overnight, except for ERα (3 nights) and caveolin-1 (2 nights). Blots were then incubated with goat anti-rabbit or mouse IgG horseradish peroxidase (1:10,000; Vector Laboratories) for 2 hours at room temperature. Bands were visualized using an enhanced chemiluminescence (ECL) kit (Thermo Fisher Scientific). Ponceau S (whole-cell) or Flotillin (biotinylation) was used as a loading control, and immunoreactive bands were normalized to obtain the percentage of protein to Ponceau S or Flotillin ratio. Bands were immediately visualized using FluorChem E and analyzed by Alphaview software (Cell Bioscience). To normalize the data as fold change from PND23 vehicle-treated for each target, the same PND23 vehicle-treated lysates were included across blots. Samples from similar treatment groups were pooled to achieve a minimum of 10 μg total protein per western blot lane.

**Table 1.**
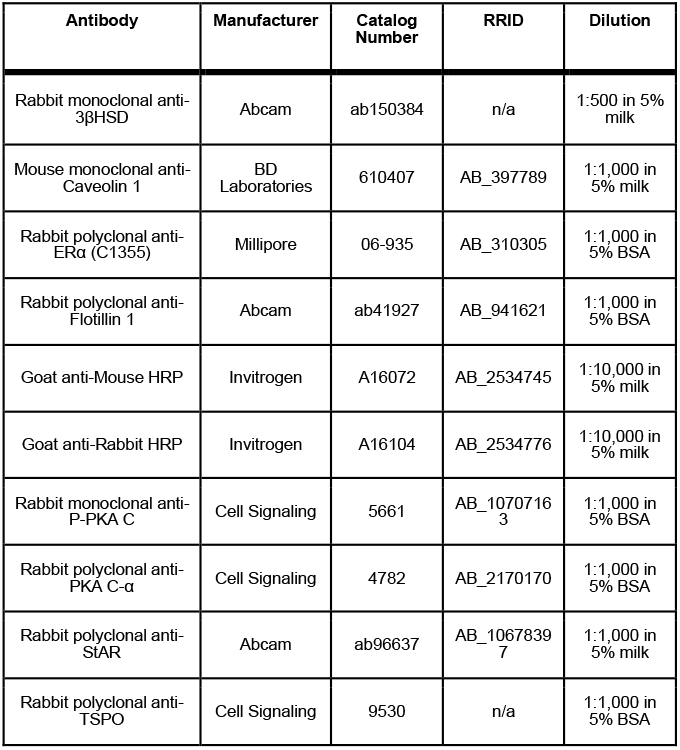
Antibody information.

### Reverse Transcription Polymerase Chain Reaction (PCR)

RNA was extracted from astrocytes using TRIzol (Invitrogen) reagent, according to manufacturer’s protocol. Following chloroform extraction of RNA and precipitation with 100% isopropanol, RNA pellet was washed with 75% ethanol/DEPC-treated water. RNA pellets dried for 10 min at room temperature and were re-suspended in DEPC-treated water. RNA concentration and quality was assessed using a spectrophotometer (NanoDrop 1000). RNA was then converted to cDNA using Superscript III First Strand Synthesis Kit (Invitrogen, cat #18080- 051). cDNA was made using Oligo dTs for Tspo, Esr1, and HSD3B1, and random hexamers for StAR using 1-2 µg RNA. The RT reaction was performed using a thermal cycler (Agilent Technologies, Stratagene Mx3000P) with the following parameters: for Oligo dTs, 50 min at 50°C and 5 min at 85°C; for random hexamers, 10 min at 25°C, 1 hr at 50°C, 10 min at 85°C. cDNA was used immediately for qPCR or stored at -20°C for ≤ 1 month.

See Table 2 for primer information. PCR reactions were run in duplicate (if using SYBR GreenER primers) or triplicate (if using TaqMan primers), and at least 3 duplicates/triplicates per condition were included in each PCR assay. No template controls (NTC; cDNA reverse transcribed without RNA) were run alongside cDNA for each primer pair in every PCR experiment. No amplification was observed in NTC wells. PCR reactions were prepared using 2 µL of cDNA in a 20 µL reaction with SYBR GreenER qPCR SuperMix Universal (Invitrogen; Carlsbad, CA) or Taqman Gene Expression Master Mix (Thermo Fisher, cat # 4369016). Conditions for amplification using SYBR GreenER primers were as follows: 2 min at 50°C (UDG incubation), 10 min at 95° C for UDG inactivation and DNA polymerase activation, then 50 cycles each (SYBR GreenER primers) or 40 cycles each (TaqMan primers) consisting of 15 sec at 95° C and 60 sec at the annealing temperature (60°C).

**Table 2.**
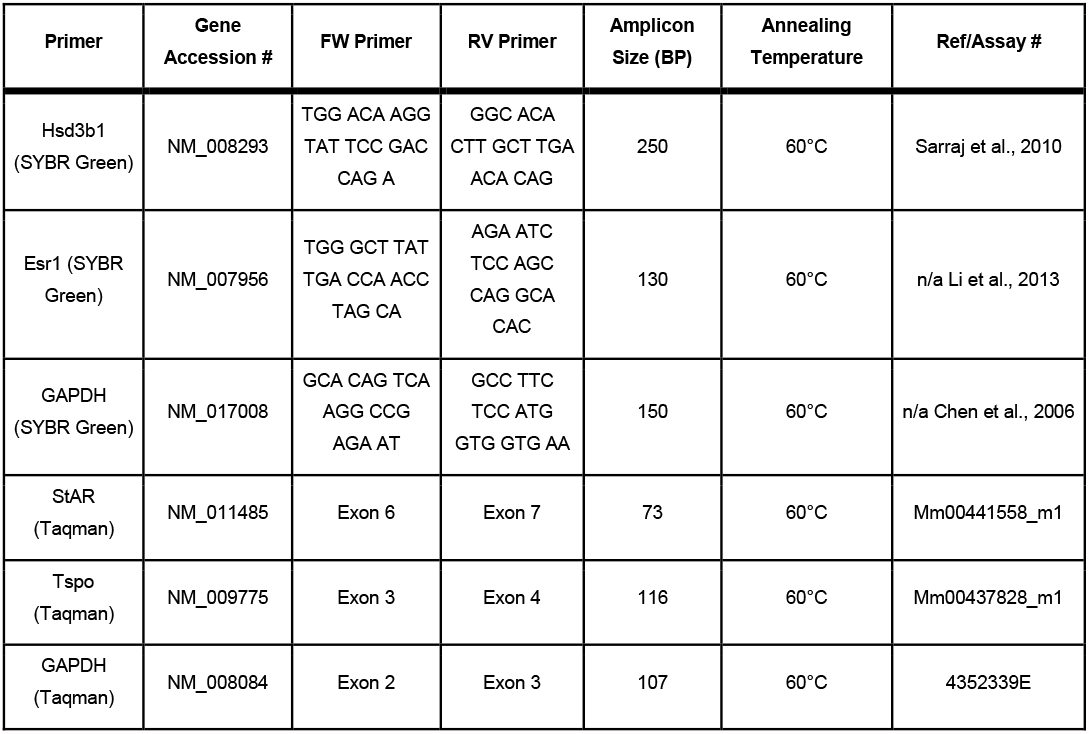
PCR Primer Information.

### Statistics

Data are presented as means ±SEM. Statistical comparisons were made using GraphPad Prism 7.01 Software (La Jolla, California). T-tests were done to determine if mRNA or peak delta F/F changes as a function of age. One-way ANOVAs with Tukey’s multiple comparisons were performed to determine if progesterone changes as a function of time. Two-way ANOVAs with Fisher’s LSD post hoc tests were performed to determine if protein levels or P4 synthesis change as a function of pubertal status and/or estrogen treatment. Statistical significance was considered if *p* ≤ 0.05.

## RESULTS

### P4 levels increased across puberty in the rostral hypothalamus of the female mouse

After treating OVX female mice with E2 (s.c.; 1 μg/20g BW; 3 hrs after lights-on), brains were collected the next day 1 hour prior to lights-out, steroids were extracted, and neuroP was measured to determine if P4 synthesis increases across pubertal development (Figure2). Two-way ANOVAs with age and hormone treatment as independent variables and progesterone (pg/ml) as the dependent variable were performed and revealed a main effect of hormone treatment, F(1, 24) = 6.22, *p* = 0.0200 and Fisher’s LSD posthoc tests indicated a post-pubertal increase in progesterone only in the anterior hypothalamus (*p* = 0.0486).

**Figure 2.**
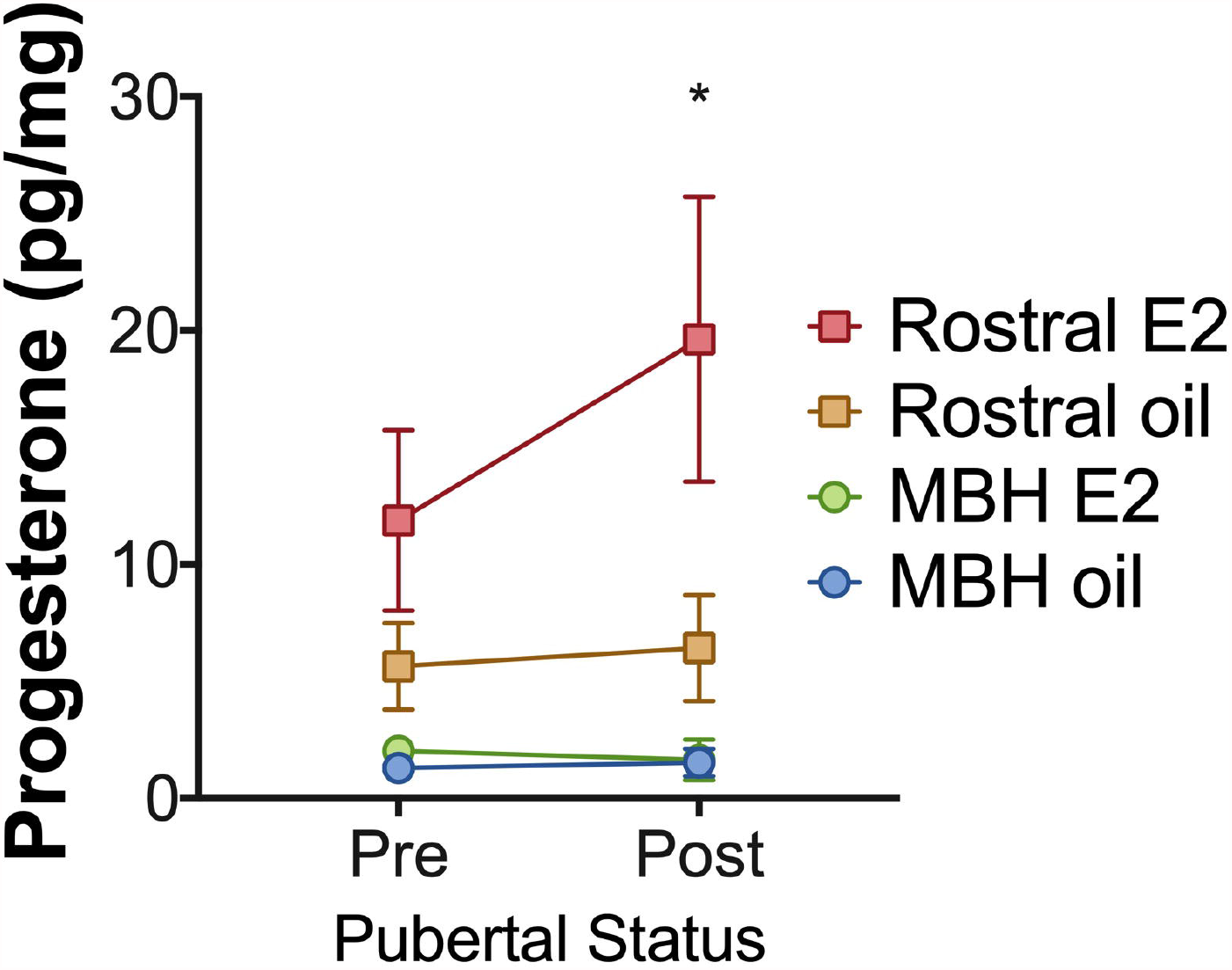
E2-induced neuroprogestone is localized to the post-pubertal rostral hypothalamus. Female mice OVX given a 1 μg E2 pellet. A week later, mice were injected with EB (s.c.; 1 μg/20g BW; 3 hrs after lights-on) and the next day brains were collected 1 hour prior to lights off (n = 6-9/group; *p = 0.0486).

### E2-induced [Ca^2+^]_i_ in hypothalamic astrocytes across puberty

To determine whether the response to E2 changes during pubertal development, prepubertal and adult female mouse hypothalamic astrocytes loaded with a calcium indicator (Fluo-4 AM) were treated with E2 and the change in fluorescence from baseline was monitored using epi- fluorescence (Figure3). Pre-pubertal hypothalamic astrocytes exhibit both an attenuated and sustained release of intracellular calcium in response to E2 treatment, compared with the rapid and transient response of these cells after puberty (Figure 3B-D). Unpaired t-tests comparing peak delta F/F indicated that adult responses to 100 pM E2 were significantly higher than the prepubertal responses to 100 pM E2 (t (20) = 7.517, *p* <0.0001; Figure 3C). Using ATP as a control for inducing intracellular calcium levels, both pre- and post-pubertal astrocytes were capable of having rapid and equally large responses (Supplementary Figure 1).

**Figure 3.**
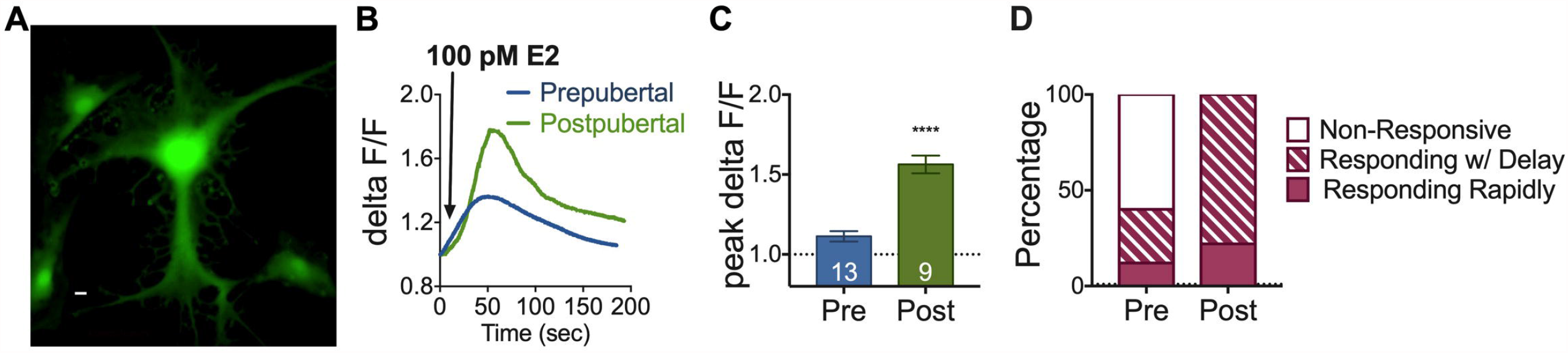
E2-induced Ca2+ dynamics change from pre- to post-puberty in hypothalamic astrocytes. (A) Fluo-4AM expression in primary hypothalamic astrocytes. (B) Representative traces of [Ca]i (delta F/F) in hypothalamic astrocytes harvested from prepubertal (PND 22) or post-pubertal (PND 60) female mice after application of 100 pM E2. (C) One way ANOVA of mean (± SEM) peak delta F/F in pre-pubertal and post-pubertal hypothalamic astrocytes reveals a significant increase in [Ca]i post-pubertal hypothalamic astrocytes compared with pre-pubertal hypothalamic astrocytes after application of 100 pM E2. (D) Percentage of cells responding to 100 pM E2. A rapid response was considered if peak delta F/F >1.2 and occurred within 50 seconds of application. A delayed response was considered if peak delta F/F >1.2 occurred at any point during application. Number of cells indicated inside bars.

### Steroidogenic proteins

Neither levels of mRNA coding for StAR, TSPO, and 3β-HSD, nor the resulting proteins differed from pre- to post-puberty (Supplementary Figure 2). Separate two-way ANOVAs with age and 10 nM E2 treatment as independent variables and fold change in protein (normalized to Ponceau S) from prepubertal vehicle-treated as the dependent variable was performed for StAR, TSPO, and 3β-HSD. StAR, TSPO, and 3β-HS protein levels did not change from pre- to post-puberty in female mouse hypothalamic astrocytes. T-tests were performed with age as the independent variable and fold change in mRNA from prepubertal astrocytes as the dependent variable revealed no significant differences in levels of StAR, TSPO, or 3β-HSD mRNA from pre- to post-puberty in female mouse hypothalamic astrocytes in vitro.

### ERα mRNA and membrane ERα

Esr1 mRNA levels in female mouse hypothalamic astrocytes harvested from prepubertal and adult female mice were compared using PCR. No changes in Esr1 mRNA were detected from pre- to post-puberty in female mouse hypothalamic astrocytes (T-test; *p* = 0.1693; Figure 4A). Surface biotinylation and western blotting revealed that the protein levels of the full-length (66kDa) ERα changed from pre- to post-puberty but not in response to 1 nM E2 treatment (Figure 4B). Two-way ANOVA with age and hormone treatment as independent variables and protein level of mERα (fold change from prepubertal vehicle-treated) as the dependent variable revealed a significant main effect of age on mERα levels (F(1, 38) = 9.35, *p* = 0.0041). There was a post-pubertal increase in mERα in female mouse hypothalamic astrocytes.

**Figure 4.**
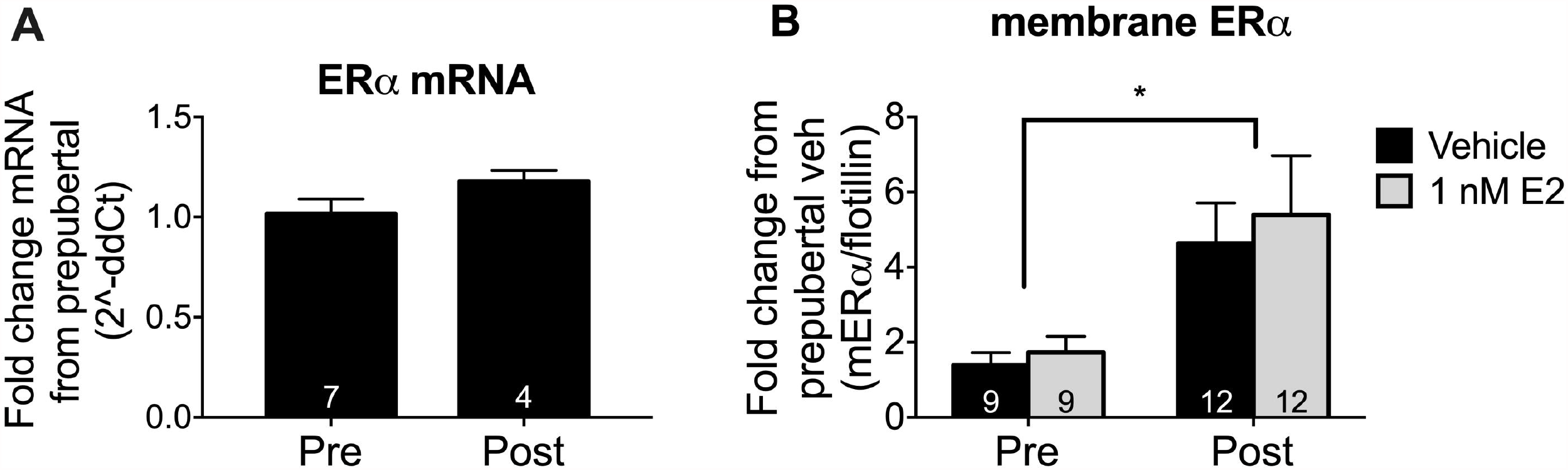
Esr1 mRNA levels are stable while mERα increase across pubertal development in female mouse hypothalamic astrocytes. A) ERα (Esr1) mRNA did not differ in hypothalamic astrocytes taken from female mice pre- or post-puberty. B) Independent of 1 nM E2-treatment, pubertal development significantly altered membrane ERα protein levels in surface biotinylated female mouse hypothalamic astrocytes. Full-length (66kDa) mERα levels increased after puberty in surface biotinylated lysates taken from female mouse hypothalamic astrocytes (ME of age, \**p* = 0.0041). Data are plotted as fold change from prepubertal vehicle mERα normalized to flotillin ± SEM. Sample size (n) for each condition is indicated within bars.

### Caveolin-1 protein

Caveolin-1 protein is required for ERα trafficking to the membrane (Meitzen and Mermelstein, 2011). Because ERα increased post-puberty, we also examined the expression of caveolin-1 protein. Whole-cell lysates of E2 or vehicle treated pre- and post-pubertal hypothalamic astrocytes were analyzed by western blotting (Figure 5A). Two-way ANOVA with age and hormone treatment as independent variables and the fold change in caveolin-1 protein (normalized to Ponceau S) from vehicle-treated prepubertal astrocytes as the dependent variable revealed a significant main effect of age (F(1, 14) = 8.486, *p* = 0.0113).

**Figure 5.**
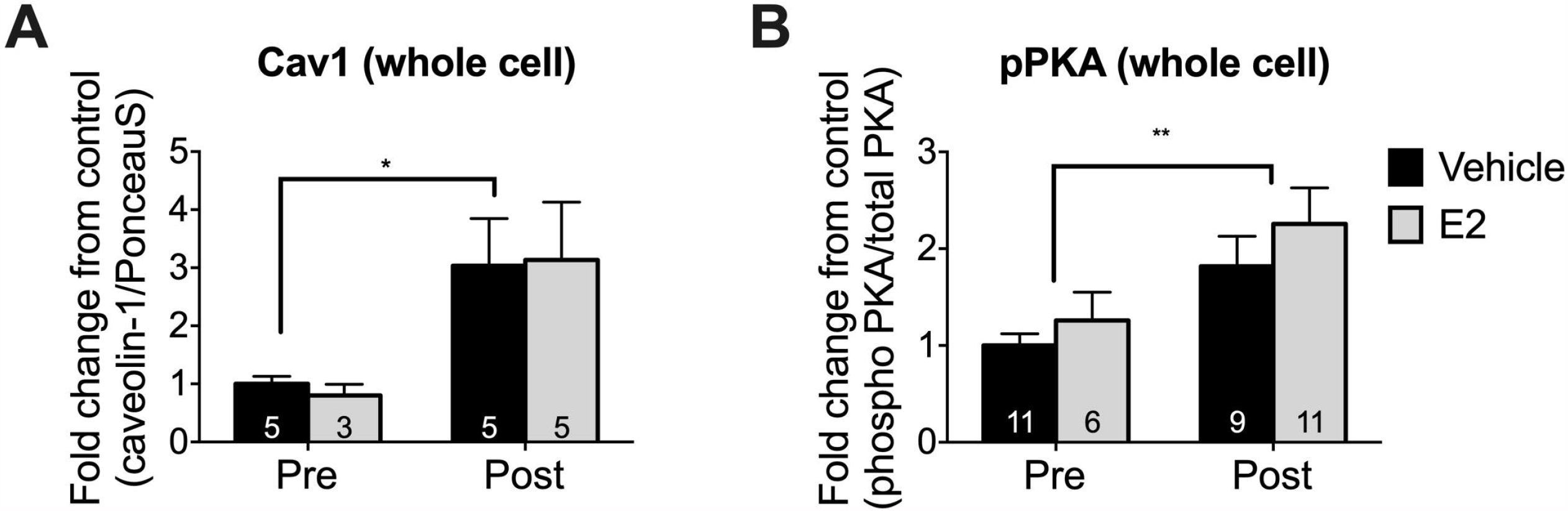
Caveolin-1 protein and PKA phosphorylation increased in adulthood in female mouse hypothalamic astrocytes. Independent of 1 nM E2-treatment, caveolin-1 protein (A) and phosphorylated PKA (B) levels significantly increased after puberty in whole-cell lysates taken from female mouse hypothalamic astrocytes (ME of age, \**p* = 0.0113; \*\**p* = 0.0052). Data are plotted as fold change in caveolin-1 or phosphorylated PKA/total PKA protein from prepubertal vehicle-treated ± SEM. Sample size (n) for each condition is indicated within bars.

### Phosphorylated PKA increased across puberty

We did not observe any changes in steroidogenic protein levels from prepuberty into adulthood, but there was a developmental increase in neuroP levels, suggesting that the activation of steroidogenic proteins was increasing. Membrane ERα signaling activates PKA to phosphorylate StAR and TSPO (11) to then increase the transport of cholesterol into the mitochondrial matrix (14), the rate-limiting step of steroidogenesis. Levels of phospho-PKA from whole-astrocyte lysates were normalized to total PKA at each developmental time point (Figure 5B). Two-way ANOVA, with age and hormone treatment as independent variables and the fold change in phospho-PKA/total PKA from prepubertal vehicle-treated as the dependent variable, revealed a significant main effect of age (F(1, 33) = 8.969, *p* = 0.0052).

## DISCUSSION

The major findings of the present study are: i) E2-induced neuroP occurs in the AVPV region of the rostral hypothalamus, but not in the mediobasal, hypothalamus; ii) this response to E2 was measured only after puberty; iii) post-pubertal changes in the steroidogenic pathway in female mouse hypothalamic astrocytes were coincident with the post-pubertal E2-induced neuroP in the rostral hypothalamus. The augmented response to E2 across pubertal development paralleled the increase of mERα levels. The increase in mERα was reflected in downstream signaling: greater PKA activation and intracellular Ca^2+^ levels. Our current understanding of the female neuroendocrine circuit that regulated ovulation implicates neuroP in augmenting E2-induced kisspeptin expression and release. Indeed, PGR signaling in AVPV kisspeptin neurons is necessary for the LH surge [Stephens, 2015; Mohr et al., 2021]. This kisspeptin, in turn, stimulates GnRH neurons and the released GnRH induces an LH surge.

The current results in female mice are congruent with earlier results in female rats that demonstrate E2-facilitated neuroP synthesis increases across puberty (15). The present findings indicated that the post-pubertal increase in E2-faciliated neuroP synthesis, happens only in the rostral hypothalamus of female mice. The mediobasal hypothalamus had the same detectable levels of progesterone, regardless of pubertal status or hormone treatment. While unlikely, an explanation for the lack of E2-induced neuroP in the MBH may be due to our E2 replacement paradigm. All mice received E2 pellets at the time of OVX, meant to mimic estrogen negative feedback, but only half received the s.c. injection of E2, meant to mimic estrogen positive feedback. It is possible that if oil-treated mice had received an oil pellet at the time of OVX, we might have observed different levels of P4 in the MBH with oil and E2 treatment; however, we designed our experiment to focus on estrogen positive feedback which requires low physiological levels of estrogen (observed in estrogen negative feedback) to prevent the downregulation of estrogen receptors and their signaling pathways.

Both pre- and post-pubertal female mouse hypothalamic astrocytes responded to E2 administration with increased [Ca^2+^]_i_, however their responses differed. Prepubertal hypothalamic astrocytes had a smaller, longer-lasting E2-induced [Ca^2+^]_i_ response. Adult hypothalamic astrocytes had a rapid, higher and more transient [Ca^2+^]_i_ response to E2. Whether these differences have functional significance remains to be characterized. When Kuo et al., (2010) compared [Ca^2+^]_i_ responses to E2 in adult male astrocytes with responses in adult female astrocytes a similar effect was observed – males had an attenuated [Ca^2+^]_i_ response. Moreover, male astrocytes, which synthesize a low level of neuroP, were not be stimulated by E2 to increase their neuroP synthesis. The response to E2 was similar in prepubertal female astrocytes and adult male astrocytes, further suggesting that adult female hypothalamic astrocytes undergo pubertal changes to allow them to synthesize neuroP. The present experiments revealed that only in post-pubertal astrocytes was PKA activated. PKA regulates the proteins needed for importing cholesterol into the mitochondrial matrix, the rate-limiting step in steroidogenesis. Thus, E2 appears to augment neuroP synthesis by increasing cholesterol transport to the mitochondria (11). These findings suggest that in pre-pubertal astrocytes, the low levels of mERα levels were not sufficient to activate intracellular signaling and that the increased trafficking of ERα to the membrane during puberty, results in adult astrocytes that can have an E2-facilitation of neuroP synthesis.

Estradiol signaling at the membrane is dependent on the mGluR transactivated by mER (reviewed in (16)). For E2- facilitated neuroP synthesis to occur, the mERα/mGluR1a complex is trafficked to the cell membrane by caveolin-1 (17) and signals through an excitatory Gq to activate PLC and the release of Ca^2+^ from intracellular stores (10). Conversely, inhibitory E2 signaling is mediated by mGluR2/3 activation of a Gi (18). Indeed, it has been shown that inhibitory E2 signaling in the hypothalamus was mediated by mGluR2/3 that was transactivated by a 52 kDa splice variant of ERα missing exon 4, the ERαΔ4 (19). In the present study, estradiol was able to elicit an attenuated [Ca^2+^]_i_ in prepubertal astrocytes, perhaps due to the low-levels of full-length mERα. In these astrocytes, E2 could be signaling through an entirely different membrane-associated receptor, GPR30, whose signaling can mediate rapid E2 effects (20). As mERα and caveolin-1 increase with development, the E2-induced synthesis of hypothalamic neuroP also increases.

We quantified mRNA and proteins levels of steroidogenic proteins to assess whether our observed developmental changes in neuroP synthesis were due to transcriptional or translational modulation. We observed no changes from pre- to post-puberty in protein levels of TSPO, StAR, or 3β-HSD, with no change seen in response to E2 treatment. TSPO’s role in steroidogenesis remains controversial, but TSPO knockouts have implicated this protein in hormone-induced steroidogenesis (21). However, the lack of effect on TSPO, StAR, or 3β-HSD suggest that the post-pubertal responsiveness to E2 is not likely attributable to up-regulation of steroidogenically relevant proteins. Given that PKA phosphorylation significantly increased post-pubertally, and that PKA phosphorylation is necessary for the *phosphorylation* of TSPO and StAR, we predict that TSPO and StAR phosphorylation increased post-pubertally, and was the likely mechanism that resulted in E2-facilation of neuroP synthesis post-pubertally.

## Conclusion

In the present study, the rostral hypothalamus responds to E2 with increased levels of neuroP but only in the post-pubertal mouse. The response to E2 in prepubertal hypothalamic astrocytes resulted in [Ca^2+^]_i_ increase but perhaps due to low amounts of full-length mERα, and caveolin-1 protein, PKA was not phosphorylated. The response to E2 in post-pubertal hypothalamic astrocytes was rapid, resulting in [Ca^2+^]_i_ flux potentially mediated by the concomitant increase in levels of full-length mERα and caveolin-1, leading to increased PKA phosphorylation. The expression of full-length mERα and caveolin-1 after puberty in hypothalamic astrocytes likely underlies the female’s distinctly post-pubertal response to E2 to produce neuroP, which is critical for ovulation.

## Acknowledgements

The authors wish to thank Carolina Thörn Perez for technical assistance during calcium imaging. This research was supported by NIH National Center for Advancing Translational Science (NCATS) UCLA CTSI grant number UL1TR001881 (MAM), and NIH Eunice Kennedy Shriver National Institute of Child Health & Human Development grant numbers HD042635 (PEM) and HD097965 (MAM).

