## Supplemental Figures 1 and 2 for "Estradiol-induced progesterone synthesis develops post-puberty in the rostral hypothalamus and coincides with post-pubertal changes in the steroidogenic pathway in female mouse hypothalamic astrocytes"

### SUPPLEMENTARY FIGURES

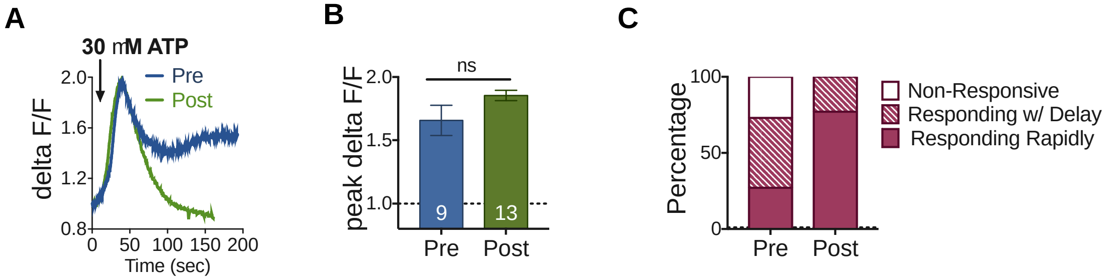


**Supplementary Figure 1. No differences in response to ATP in pre-and post-pubertal hypothalamic astrocytes** (A) Representative traces of [Ca]i (delta F/F) in hypothalamic astrocytes harvested from postnatal day 22 (pre-pubertal) or 60 (post-pubertal) female mice after application of 30 uM ATP. (B) One way ANOVA of mean (± SEM) peak delta F/F in pre-pubertal and post-pubertal hypothalamic astrocytes indicates no significant differences in [Ca]i post-pubertal hypothalamic astrocytes compared with pre-pubertal hypothalamic astrocytes after application of 30 uM ATP. (C) Percentage of cells responding to 30 uM. A rapid response was considered if peak delta F/F >1.2 and occurred within 50 seconds of application. A delayed response was considered if peak delta F/F >1.2 occurred at any point during application.

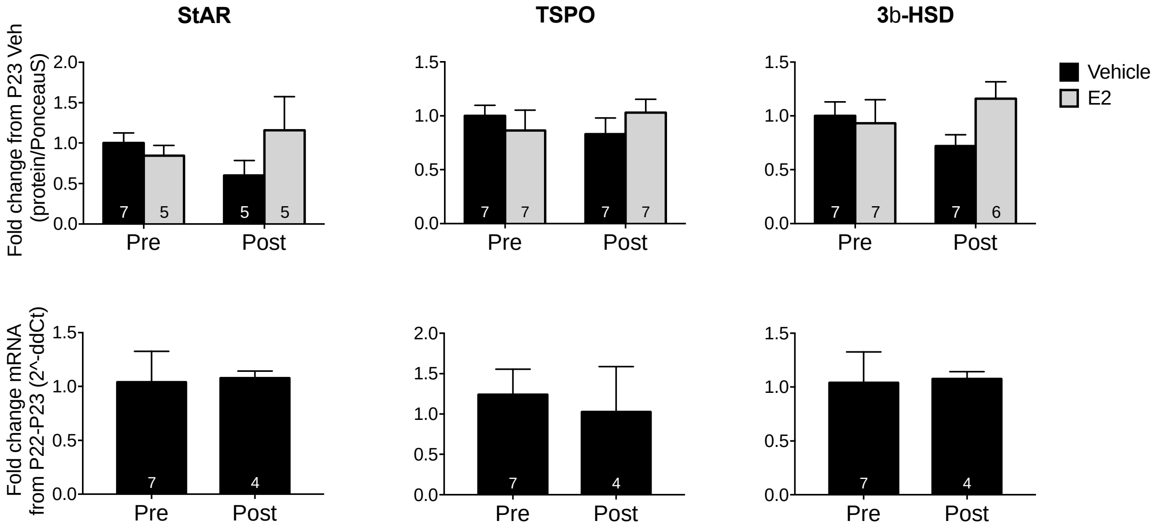


Supplementary Figure 2. No differences in StAR, TSPO, or 3b-HSD protein or mRNA from pre- to post-puberty in female mouse hypothalamic astrocytes. Two-way ANOVAs with protein and age as independent variables and protein expression as the dependent variable indicated no main effects of age or hormone treatment, and no interactions on protein expression. T-tests with age as the dependent variable and mRNA as the independent variable indicate no significant differences in mRNA between pre- and post-puberty in female mouse hypothalamic astrocytes. Significance set at $p < 0.05$, sample size (n) of different conditions indicated inside bars.
